# Positive Transfer of the Whisper Speech Transformer to Human and Animal Voice Activity Detection

**DOI:** 10.1101/2023.09.30.560270

**Authors:** Nianlong Gu, Kanghwi Lee, Maris Basha, Sumit Kumar Ram, Guanghao You, Richard H. R. Hahnloser

## Abstract

This paper introduces WhisperSeg, utilizing the Whisper Transformer pre-trained for Automatic Speech Recognition (ASR) for human and animal Voice Activity Detection (VAD). Contrary to traditional methods that detect human voice or animal vocalizations from a short audio frame and rely on careful threshold selection, WhisperSeg processes entire spectrograms of long audio and generates plain text representations of onset, offset, and type of voice activity. Processing a longer audio context with a larger network greatly improves detection accuracy from few labeled examples. We further demonstrate a positive transfer of detection performance to new animal species, making our approach viable in the data-scarce multi-species setting.^1^

## 1. INTRODUCTION

Voice Activity Detection (VAD) is the task of identifying the onset, offset, and type of speech in extended audio recordings, foundational in speech recognition [1], speaker diarization [2], and speech analysis tasks that require precise speech segmentations at word or utterance levels [3, 4, 5]. This task extends to the domains of animal communication [6] and vocal learning research [7], where vocalizations must be segmented against background noises [8, 9]. Given the intrinsic similarities between human and animal VAD, there is potential to develop a universal vocal segmentation tool that can effectively handle a wide range of species. Such a tool would streamline segmentation efforts and potentially contribute to comparative research on vocal communication.

Recent approaches to human and animal VAD typically use frame-by-frame classification strategies [10, 11, 12], where an audio spectrogram is divided into fixed-length frames (e.g., 20 to 30 columns within the spectrogram). A neural network such as a Convolutional Neural Network (CNN) then processes each frame to produce a hidden state. This state is subsequently processed by a recurrent neural network, like an LSTM, generating a final vector representation for each frame that is used for frame-wise classification. Depending on a selected threshold (e.g., 0.5), each frame is classified as non-voice (0) or voice (1), with additional classes to resolve vocalization type if necessary. Analyzing the resulting sequence of classifications allows the extraction of vocal activity details such as onset, offset, and type.

Traditional frame-wise classification approaches come with inherent challenges: 1) The classification of each frame requires an empirically chosen decision threshold. This threshold, applied universally to all frames, can falter in real-world recordings where nonstationary environmental noises and varying recording quality introduce substantial audio heterogeneity. 2) Audio frames may lack sufficient contextual information for accurately detecting vocal activity. Particularly with human speech, discerning between a pause within a structured vocal unit (e.g. an utterance) and true silence can be difficult without a broader context.

The recent Whisper Transformer is a speech-to-text encoder-decoder model trained on massive data [13]. In principle, Whisper overcomes the above challenges due to its threshold-free textual output and long input spectrograms with rich contexts. While prior studies have leveraged Whisper for human speech timestamp prediction [13, 14], its broader application remains relatively uncharted. This work investigates the transfer of Whisper to universal voice detection, targeting both human and animal vocalizations, elevating its potential as an all-encompassing VAD solution.

## 2. METHOD

### Our methodology comprises the following steps

#### Preprocessing and Adaptation

From an audio clip with sampling rate *f*_*s*_ and duration *D*, we compute the log-Mel spectrogram using 80 Mel bands within a species-specific frequency range (Table 1) and hop length *L*_hop_ between FFT windows, to yield a spectrogram with exactly 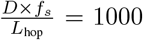 columns. *D* and *L*_hop_ are tailored for each species to account for varying utterance durations, such that one spectrogram contains a similar number of vocal segments across species (e.g., up to 30 segments per audio clip) and each segment spans a consistent number of spectrogram columns (e.g., 20 to 300 columns). By reducing inter-species differences in the input spectrogram, we can fine-tune a universal WhisperSeg model to detect onsets and offsets using spectrogram column indices rather than raw timestamps. This strategy obviates the need to add or adapt timestamping tokens for individual species, promoting a uniform segmentation procedure irrespective of the utterance duration and vocalizing species.

**Table 1:** Statistics for the multi-species dataset and settings when generating log-Mel spectrograms.

| Dataset | No. segments | | Segment length (s) | | SNR (dB) | No. segment types | audio clip duration $D$ (s) | sampling rate $f_s$ (kHz) | nfft | hop length $L_{hop}$ | minimum frequency (kHz) | maximum frequency (kHz) | WhisperSeg time resolution (s) | $\epsilon_{vote}$ (s) |
| --- | --- | --- | --- | --- | --- | --- | --- | --- | --- | --- | --- | --- | --- | --- |
|  | train | test | train | test |  |  |  |  |  |  |  |  |  |  |
| Zebra Finch (adults) | 36,226 | 3,909 | $0.15 \pm 0.14$ | $0.15 \pm 0.14$ | 20.9 | 1 | 2.5 | 32 | 512 | 80 | 0 | 16 | 0.005 | 0.02 |
| Zebra Finch (juveniles) | 12,312 | 980 | $0.13 \pm 0.07$ | $0.13 \pm 0.07$ | 11.3 | 1 | 2.5 | 32 | 512 | 80 | 0 | 16 | 0.005 | 0.02 |
| Bengalese Finch | 28,562 | 2,074 | $0.07 \pm 0.03$ | $0.07 \pm 0.03$ | 9.0 | 1 | 2.5 | 32 | 512 | 80 | 0 | 16 | 0.005 | 0.02 |
| Marmoset | 2,535 | 702 | $0.13 \pm 0.14$ | $0.11 \pm 0.12$ | 6.0 | 4 | 2.5 | 48 | 1024 | 120 | 0 | 24 | 0.005 | 0.02 |
| Mouse | 1,807 | 229 | $0.06 \pm 0.05$ | $0.05 \pm 0.04$ | 17.5 | 1 | 0.5 | 300 | 4096 | 150 | 35 | 150 | 0.001 | 0.02 |
| Human (AVA Speech) | 21,958 | 93 | $3.39 \pm 5.17$ | $3.47 \pm 3.95$ | 4.3 | 1 | 10 | 16 | 512 | 160 | 0 | 8 | 0.02 | 0.2 |

During conversion of onset and offset timestamps *t* into column indices for fine-tuning, we round 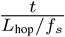 to the nearest even number. This aligns with Whisper’s native time resolution, which is half that of the spectrogram due to the 2x down-sampling in Whisper’s convolution layers (Figure 1), which effectively halves the input size to Whisper encoder [13].

**Fig. 1:**
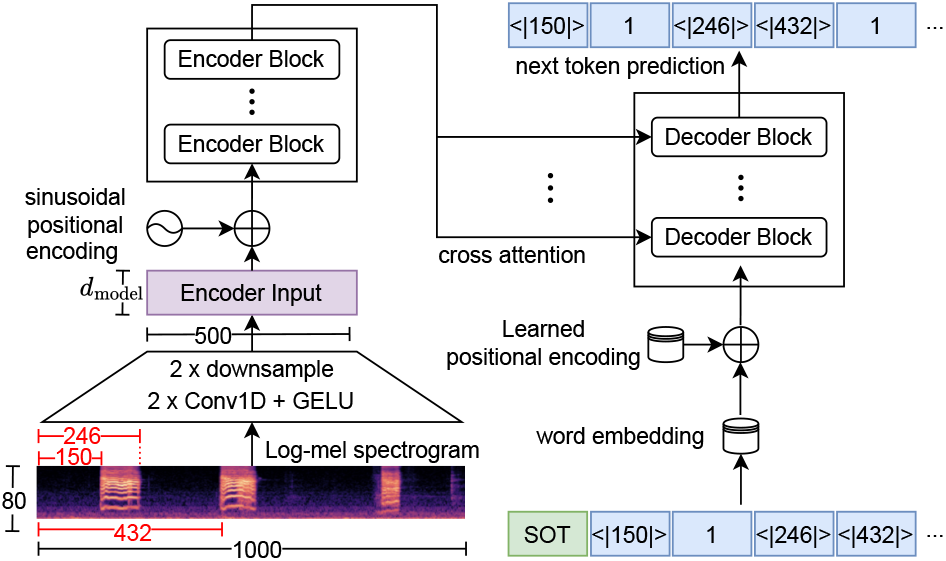
WhipserSeg is trained to generate transcripts of vocal segments with onsets and offsets specified as column indices.

#### Model Fine-tuning

Utilizing the spectrograms and column indices obtained, we finetune Whisper to generate text sequences representing vocal activity. In the example in Figure 1, the target output is: `SOT<|150|>1<|246|><|432|>1 <|518|><|810|>1<|854|>`, where SOT marks the start of decoding. The subsequent `<|150|>1<|246|>` indicates the onset of a vocal segment type (ID) 1 spanning from the 150th to the 246th spectrogram column.

#### Segment Extraction and Temporal Mapping

Segments are extracted from generated sequences using regular expressions. The onset and offset column indices are then mapped to timestamps relative to the beginning of the audio clip by multiplying the column index with the spectrogram’s temporal resolution *L*_hop_*/f*_*s*_. The extracted segment type IDs are converted to the original segment type using a mapping table.

#### Segmentation of Extended Audios

To process extended sound files, we divide them into fixed-length clips (of duration *D* outlined in Table 1) and apply WhisperSeg to each. Consecutive segments from adjacent clips of the same type are concatenated (Figure 2a). However, the dividing point of sound clips may affect accuracy, especially when it inadvertently bisects a vocalization. To counter this effect, we process a sound file 3 times, each time with a distinct division point that we incrementally shift by 1/3 of the clip duration, producing for each sound file three segmentation variants. To obtain final segments, we use a majority voting strategy: a segment is considered valid if, in another variant, there is a counterpart segment with a mean absolute difference (of both onsets and offsets) below a predetermined threshold *ϵ*_vote_. The final onset and offset are then defined as the averages across all valid segments. Segments without support from another co-valid segment are discarded (Figure 2b).

**Fig. 2:**
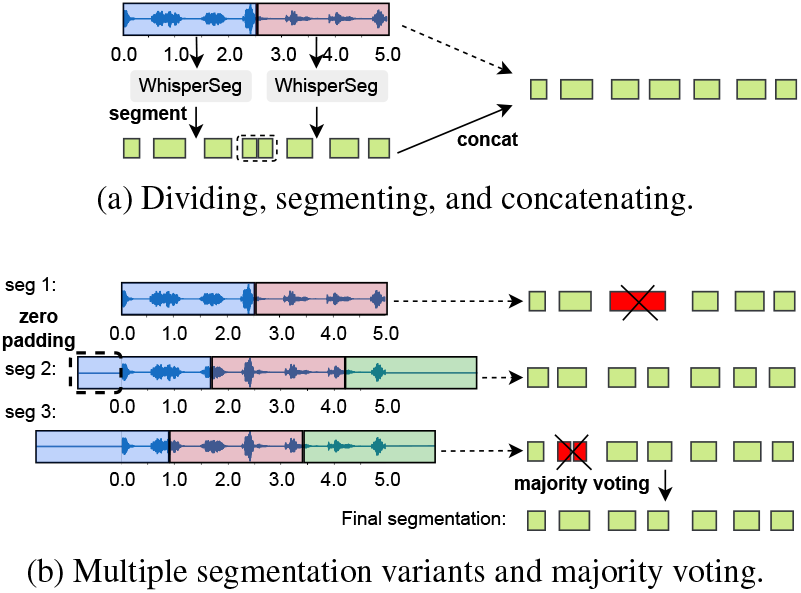
Strategies for segmenting extended audios.

## 3. DATASET

We evaluate WhisperSeg on five species datasets detailed in Table 1. The zebra finch [15] and Bengalese finch [12, 16] datasets represent bird recordings in controlled, quiet environments. The marmoset [12, 17] and mouse [12, 18] datasets cover mammalian sounds; notably, mouse vocalizations occur in the ultrasonic domain, while marmoset recordings feature elevated noise levels, measured by signal-to-noise power ratio (SNR), due to their origin from a housing room with multiple animals. For human speech, the AVA-Speech dataset, derived from film audio [19, 10], captures diverse noisy conditions. These datasets help assess WhisperSeg’s performance over various species, frequencies, and noise levels.

Table 1 shows the multi-species dataset statistics and settings for log-Mel spectrogram computation. For mice vocalizations, due to their ultrasonic nature, the log-Mel spectrogram contains 80 Mel bands starting at a minimum frequency of 35 kHz. The voting threshold *ϵ*_vote_ is empirically determined based on the segment length and WhisperSeg’s time resolution. To prevent naming conflicts, we prefixed each vocalization type with a token representative of the respective species. This prefixing ensures that each vocalization type from every species is given a unique type ID. As a result, the WhisperSeg trained on multi-species datasets can discern both the type of vocalization and the species producing it.

## 4. EXPERIMENTS AND DISCUSSION

We fine-tuned WhisperSeg for next-token prediction using cross-entropy loss. We used two model sizes: a base version with 74M parameters (whisper-base) and a larger 1.55B parameter variant (whisper-large) [13]. We explored training species-specific WhipserSeg models and training a single WhipserSeg on all five species. Training parameters encompassed three epochs, a 3e-6 learning rate introduced with a linear warm-up over 100 model update steps followed by a linear decay to zero, and a batch size of 4. We augmented the training process by randomly choosing spectrogram onsets during training, to increase the number of positions at which vocalizations are sampled. Subsequent sections detail baselines, evaluation metrics, and performance analysis across species, assessing WhisperSeg’s adaptability.

### Baselines

We compared WhisperSeg with 1) VoxSeg [10], which uses a time-distributed CNN to convolve 25-column wide frames, shifting by 2 columns each time. Given a padded 1000-column spectrogram, this results in 500 hidden vectors, which are subsequently processed by a Bi-LSTM to produce 500 encoded outputs for frame-wise classification (e.g., voice vs noise); 2) DAS [12], which introduces a learned FFT block to transform audio samples into features that are subsequently processed by Temporal Convolutional Networks (TCNs) [20]; 3) MLPSeg, an additional simple baseline we implemented that classifies a 25-column wide spectrogram frame through four fully connected layers. While these models diverge in contextual scope, they commonly predict labels for brief time frames, from which they derive onset, offset, and vocalization types. Contrarily, WhisperSeg directly infers these parameters through token generation without explicitly resorting to a serial computation at a high temporal resolution.

### Evaluation Metrics

We evaluated segmentation performance using segment-wise (F1_seg_) and frame-wise (F1_frame_) F1 scores. F1_seg_ aligns model predictions with ground truth, considering a tolerance of segment onset, offset, and type consistency of 200 ms for humans and 10 ms for all other species. From this alignment, we compute numbers of true positive (TP), false positive (FP), and false negative (FN) test samples to obtain the F1 score (Figure 3). F1_frame_ divides time into short bins of 10 ms for humans and 1 ms for other species and compares predicted and actual vocal segment types bin-by-bin. Notably, F1_seg_ is more stringent, prioritizing segment integrity and penalizing minor segmentation errors.

**Fig. 3:**
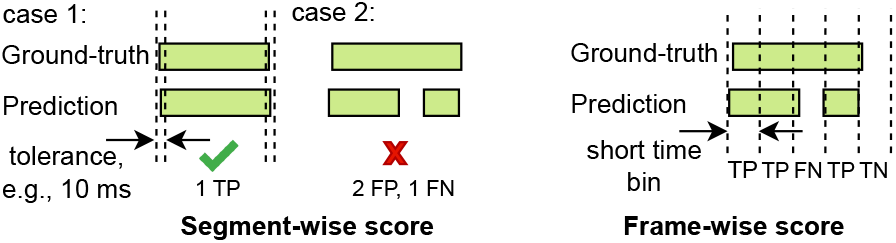
Illustrations of evaluation metrics.

## Results and Discussion

WhisperSeg outperformed base-line models in both F1_seg_ and F1_frame_ on human and animal datasets, especially on noisy datasets from Marmosets and Humans. While baseline models achieve commendable F1_frame_ scores, they falter on F1_seg_ scores under noisy conditions, often producing fragmented predictions or brief false positives. In contrast, WhisperSeg’s performance is stable on both metrics, emphasizing its accurate onset and offset predictions and strong resilience to noise. Notably, a universal WhisperSeg model (WSeg-large-MS), trained across all species, matches the performance of species-specific models (Table 2), highlighting its versatile generalization capability. By exception, the fine-tuning of a given WhisperSeg model on animal datasets provides no clear advantage for human VAD, which could be either due to the intricacies of the human data or the lack of pre-training on animal data.

**Table 2:** Comparison of animal sound segmentation and VAD performance. “WSeg-large-S3” is the large WhisperSeg variant employing a 3-variant majority voting strategy (Figure 2b), while “S1” corresponds to a simple concatenation of (1-variant) segments as shown in Figure 2a. “MS” denotes a universal model trained across all species with species-specific test scores. For the zebra finch dataset, we combined the training datasets of both adults and juveniles but presented test results separately for each group. Except for DAS, models were trained and tested 5 times, with average test scores reported.

| Model | No. parameters | Zebra Finch (adults) |  | Zebra Finch (juveniles) |  | Bengalese Finch |  | Marmoset |  | Mouse |  | Human (AVA Speech) |  |
| --- | --- | --- | --- | --- | --- | --- | --- | --- | --- | --- | --- | --- | --- |
|  |  | F1<sub>seg</sub> | F1<sub>frame</sub> | F1<sub>seg</sub> | F1<sub>frame</sub> | F1<sub>seg</sub> | F1<sub>frame</sub> | F1<sub>seg</sub> | F1<sub>frame</sub> | F1<sub>seg</sub> | F1<sub>frame</sub> | F1<sub>seg</sub> | F1<sub>frame</sub> |
| MLPSeg | 554k | 0.9601 | 0.9791 | 0.7403 | 0.9374 | 0.8895 | 0.9341 | 0.0752 | 0.1853 | 0.8352 | 0.9033 | 0.0513 | 0.8173 |
| DAS | 180k | 0.9797 | 0.9883 | 0.8489 | 0.9685 | 0.8977 | 0.9564 | 0.3026 | 0.7120 | 0.9536 | 0.9687 | 0.0461 | 0.8218 |
| VoxSeg | 619k | 0.9894 | 0.9830 | 0.9171 | 0.9723 | 0.9187 | 0.9450 | 0.1534 | 0.8106 | 0.9491 | 0.9769 | 0.2837 | 0.9300 |
| WSeg-base-S1 | 74M | 0.9940 | 0.9878 | 0.9130 | 0.9776 | 0.9710 | 0.9656 | 0.6651 | 0.7738 | 0.9190 | 0.9613 | 0.3694 | 0.9311 |
| WSeg-base-S3 | 74M | 0.9957 | 0.9901 | 0.9543 | 0.9793 | 0.9741 | 0.9737 | 0.7265 | 0.8123 | 0.9332 | 0.9658 | 0.3968 | 0.5980 |
| WSeg-large-S1 | 1550M | 0.9945 | 0.9883 | 0.9345 | 0.9800 | 0.9793 | 0.9723 | 0.7744 | 0.8573 | 0.9398 | 0.9798 | 0.3730 | 0.9366 |
| WSeg-large-S3 | 1550M | 0.9962 | **0.9905** | **0.9583** | **0.9830** | **0.9819** | **0.9781** | 0.7988 | **0.8742** | 0.9437 | 0.9801 | **0.4195** | 0.7184 |
| WSeg-large-MS-S1 | 1550M | 0.9942 | 0.9882 | 0.9364 | 0.9796 | 0.9766 | 0.9695 | 0.7933 | 0.8662 | 0.9437 | 0.9810 | 0.3797 | 0.9349 |
| WSeg-large-MS-S3 | 1550M | **0.9966** | **0.9905** | 0.9551 | 0.9827 | 0.9789 | 0.9760 | **0.8095** | 0.8741 | **0.9545** | **0.9813** | 0.3995 | 0.6386 |
| Whisper-Large-Framewise | 1550M | 0.9773 | 0.9856 | 0.8353 | 0.9740 | 0.9295 | 0.9654 | 0.1482 | 0.5336 | 0.8172 | 0.9246 | 0.3593 | **0.9505** |

WhisperSeg’s superiority arises from its transformer architecture and numerous parameters, excelling particularly in noisy environments. In addition, when compared with a frame-wise Whisper solution, where both the encoder and decoder of Whisper receive the spectrogram and the decoder’s output is used for frame-wise classification, the “Whisper-Large-Framewise” model underperformed compared to “WSeg-Large-S1”, Table 2. This suggests that the token-based approach of WhisperSeg is a key factor in its superior performance, rather than just its large model size.

We observed that longer input audio contexts significantly enhance vocal segmentation accuracy. Models with extended audio contexts, such as VoxSeg and WhisperSeg, consistently outperformed those with shorter contexts like MLPSeg, especially in low signal-to-noise ratio (SNR) scenarios. In further experiments with WhisperSeg using varying input spectrogram lengths, we noted consistent improvements in performance. Increasing the context lengths from 100, 200, to 1000 columns resulted in a steady rise in F1_seg_ scores: from 0.081, 0.222, to 0.380 in humans, and from 0.712, 0.832, to 0.944 in mice. In other species, the improvements were also exhibited but less substantial. These findings underscore the benefits of longer audio contexts for WhisperSeg’s performance.

In addition, the majority voting approach enhances F1_seg_ scores when comparing WSeg-xxx-S3 to WSeg-xxx-S1 (Table 2). A drop in F1_frame_ for the Human dataset after majority voting could be due to the large variations in segment duration, which influence predictions and may lead to segment exclusion during voting.

We examined WhisperSeg’s adaptability to new datasets and its reliance on training set size by fine-tuning WhisperSeg on a new species, starting with small subsets (50 to 1000 segments) and eventually using the full set. Notably, when selecting a zebra finch training subset of *N* segments, we ensured an even split between adult and juvenile vocal segments to enforce an age balance in training examples. We trained three models on the new species: a) VoxSeg, b) WhisperSeg-large initialized from a pretrained ASR checkpoint [13] with tri-variant majority voting (WS-L-S3), and c) WhisperSeg-large pretrained on multi-species data (excluding the new species data currently being tested) with majority voting (WS-L-MS-S3, Figure 4).

**Fig. 4:**
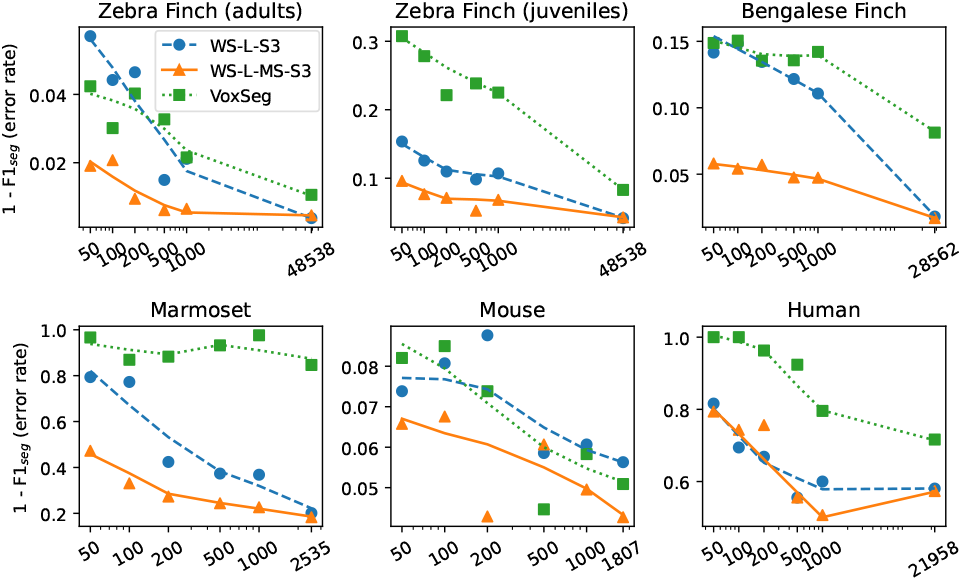
Comparison of performance on the complete test dataset of a given new species. The x-axis shows the training subset size of the new species (in segment count), with the rightmost point representing the full training set size. The y-axis depicts 1 − F1_seg_ as an error rate measure.

The multi-species WhisperSeg (WS-L-MS-S3) consistently outperformed baselines, even with minimal training data. Notably, by leveraging a mere 50 segments from a novel species, WS-L-MS-S3 achieves an error rate reduction of over 50% when compared with baseline models on zebra finch, Bengalese finch, and marmoset datasets. On zebra finch and marmoset datasets, WS-L-MS-S3 converges to performance levels seen in full-size training but only requires 200 segments, demonstrating its efficiency in rapid adaptability. This evidence underscores the profound effectiveness of our proposed multi-species Whisper-based segmenter, showcasing its promise in real-world applications where data from unfamiliar individuals or species is introduced.

## 5. CONCLUSION

In this study, we successfully adapted the Whisper Transformer, originally designed for human ASR, to Voice Activity Detection (VAD) for humans and animals. Our strategy leverages the prediction of onset and offset timestamps, expressed as column indices within fixed-size input spectrograms. This spectrogram-level preprocessing facilitates the universal application of the Whisper model across diverse animal species. Empirical results underscore a consistent superiority across all tested species and emphasize the advantages of crafting a multi-species vocalization segmenter with enhanced generalization and adaptation capabilities.

## Acknowledgements

We acknowledge support from the Swiss National Science Foundation (grant 31003A 182638) and the NCCR Evolving Language, Swiss National Science Foundation Agreement No. 51NF40 180888. We also thank the anonymous reviewers for their useful comments.

## Footnotes

1 Our code and data are at https://github.com/nianlonggu/WhisperSeg

## Notes

https://github.com/nianlonggu/WhisperSeg

### Competing Interest Statement

The authors have declared no competing interest.

### Summary of Updates

Table 3 removed; Experiment results of more baseline models added in Section 4; Acknowledgments added.

## REFERENCES

[1] Nitin N Lokhande, Navnath S Nehe, and Pratap S Vikhe, “Voice activity detection algorithm for speech recognition applications,” in IJCA Proceedings on International Conference in Computational Intelligence (IC-CIA2012), vol. iccia, 2012, vol. 6, pp. 1–4.

[2] Ivan Medennikov, Maxim Korenevsky, Tatiana Prisyach, Yuri Khokhlov, Mariya Korenevskaya, Ivan Sorokin, Tatiana Timofeeva, Anton Mitrofanov, Andrei Andrusenko, Ivan Podluzhny, et al., “Target-speaker voice activity detection: a novel approach for multispeaker diarization in a dinner party scenario,” arXiv preprint 2005.07272, 2020.

[3] Manuel Milling, Alice Baird, Katrin D Bartl-Pokorny, Shuo Liu, Alyssa M Alcorn, Jie Shen, Teresa Tavassoli, Eloise Ainger, Elizabeth Pellicano, Maja Pantic, et al., “Evaluating the impact of voice activity detection on speech emotion recognition for autistic children,” Frontiers in Computer Science, vol. 4, pp. 837269, 2022.

[4] Hannah Sarvasy, Jaydene Elvin, Weicong Li, and Paola Escudero, “An acoustic analysis of nungon vowels in child-versus adult-directed speech,” in Proceedings of the 19th International Congress of Phonetic Sciences Melbourne, 2019, pp. 3155–3159.

[5] Frank Seifart, Jan Strunk, Swintha Danielsen, Iren Hartmann, Brigitte Pakendorf, Søren Wichmann, Alena Witzlack-Makarevich, Nivja H de Jong, and Balthasar Bickel, “Nouns slow down speech across structurally and culturally diverse languages,” Proceedings of the National Academy of Sciences, vol. 115, no. 22, pp. 5720–5725, 2018.

[6] Lisa F Gill, Wolfgang Goymann, Andries Ter Maat, and Manfred Gahr, “Patterns of call communication between group-housed zebra finches change during the breeding cycle,” Elife, vol. 4, pp. e07770, 2015.

[7] Sepp Kollmorgen, Richard HR Hahnloser, and Valerio Mante, “Nearest neighbours reveal fast and slow components of motor learning,” Nature, vol. 577, no. 7791, pp. 526–530, 2020.

[8] Thomas Colligan, Kayla Irish, Douglas J. Emlen, and Travis J. Wheeler, “Disco: A deep learning ensemble for uncertainty-aware segmentation of acoustic signals,” bioRxiv, 2023.

[9] Zifan Jiang, Adrian Soldati, Isaac Schamberg, Adriano R Lameira, and Steven Moran, “Automatic sound event detection and classification of great ape calls using neural networks,” arXiv preprint 2301.02214, 2023.

[10] Nicholas Wilkinson and Thomas Niesler, “A hybrid cnn-bilstm voice activity detector,” in ICASSP 2021 - 2021 IEEE International Conference on Acoustics, Speech and Signal Processing (ICASSP), 2021, pp. 6803–6807.

[11] Yarden Cohen, David Aaron Nicholson, Alexa Sanchioni, Emily K Mallaber, Viktoriya Skidanova, and Timothy J Gardner, “Automated annotation of birdsong with a neural network that segments spectrograms,” eLife, vol. 11, pp. e63853, jan 2022.

[12] Elsa Steinfath, Adrian Palacios-Munõz, Julian R Rottschäfer, Deniz Yuezak, and Jan Clemens, “Fast and accurate annotation of acoustic signals with deep neural networks,” eLife, vol. 10, pp. e68837, nov 2021.

[13] Alec Radford, Jong Wook Kim, Tao Xu, Greg Brockman, Christine McLeavey, and Ilya Sutskever, “Robust speech recognition via large-scale weak supervision,” 2022.

[14] Jérôme Louradour, “whisper-timestamped,” https://github.com/linto-ai/ whisper-timestamped, 2023.

[15] Tomas Tomka, Xinyu Hao, Aoxue Miao, Kanghwi Lee, Maris Basha, Stefan Reimann, Anja T Zai, and Richard Hahnloser, “Benchmarking nearest neighbor retrieval of zebra finch vocalizations across development,” bioRxiv, 2023.

[16] David Nicholson, Jonah E. Queen, and Samuel J. Sober, “Bengalese Finch song repository,” 5 2021.

[17] Rogier Landman, Jitendra Sharma, Julia B Hyman, Adrian Fanucci-Kiss, Olivia Meisner, Shivangi Parmar, Guoping Feng, and Robert Desimone, “Close-range vocal interaction in the common marmoset (callithrix jacchus),” Plos one, vol. 15, no. 4, pp. e0227392, 2020.

[18] B. (Bernhard) Englitz, M.A.J. van (Marcel) Gerven, Paul Watkins, Alexander Ivanenko, and Kurt Hammer-schmidt, “Classifying sex and strain from mouse ultrasonic vocalizations using deep learning,” 2020.

[19] Sourish Chaudhuri, Joseph Roth, Dan Ellis, Andrew C. Gallagher, Liat Kaver, Radhika Marvin, Caroline Pantofaru, Nathan Christopher Reale, Loretta Guarino Reid, Kevin Wilson, and Zhonghua Xi, “Ava-speech: A densely labeled dataset of speech activity in movies,” in Proceedings of Interspeech, 2018, 2018.

[20] Shaojie Bai, J. Zico Kolter, and Vladlen Koltun, “An empirical evaluation of generic convolutional and recurrent networks for sequence modeling,” 2018.

